# Metabotropic purinergic receptor profiles and calcium signalling in primary mice myoblasts differ depending on their muscle origin and are altered in cells with mutated dystrophin gene (mdx mice)

**DOI:** 10.1101/2022.02.16.478175

**Authors:** Justyna Róg, Aleksandra Oksiejuk, Dariusz C. Górecki, Krzysztof Zabłocki

## Abstract

Mortality of Duchenne Muscular Dystrophy (DMD) is a direct consequence of progressive wasting of muscle fibres leading to skeletal muscle deterioration and cardiomyopathy. However pathophysiological effects of mutations in the dystrophin encoding gene, which result in improper muscle maturation are detectable in muscle precursor cells which do not express dystrophin gene at the protein level because of early stage of differentiation thus irrespectively of changes in dystrophin-encoding gene. Among these abnormalities elevated activity of P2X7 receptors and increased store-operated calcium entry, have been identified in mdx mouse myoblasts. Moreover the increased response of immortalized mdx myoblasts to agonists activating metabotropic purinergic receptors was characterised. Experiments on immortalized myoblasts do not allow indicating potentially specific effects of mdx mutation on cells derived from particular muscles. Moreover an immortalization of cells itself may influence their metabolism in poorly defined way. Therefore here the metabotropic response of primary myoblasts derived from various muscles of normal and mdx mice to nucleotide stimulation has been investigated. Transcript and protein level of P2Y receptors, sensitivity to antagonist, and cellular localization clearly indicate P2RY2 as the most affected in mdx myoblasts. This meets our previous conclusion drawn from experiments with immortalized cells. However a pattern of expression and activity of P2Y receptors among myoblasts derived from four muscles differ. Also cellular levels of some other proteins belonging to the “calcium signalling toolkit” differ in myoblasts from various muscle and are differently changed due to mdx mutation. Finally, these results complement and strongly support previously formulated conclusion that phenotypic effects of DMD emerge as early as in undifferentiated muscle and therefore traditional understanding of DMD pathogenesis needs re-evaluation.

## Introduction

Duchenne muscular dystrophy (DMD) is a leading neuromuscular disease which ultimately leads to severe disability and premature death due to damaged diaphragm and accessory respiratory muscles and/or cardiac complications [Yiu et al. 2015]. Moreover, DMD also affects nervous system leading to reduced cognitive capability of patients. DMD is a consequence of out-of-frame mutations in dystrophin encoding gene located on the chromosome X. Thus, it almost exclusively affects boys with a prevalence 1 per 5000 live births Crisafulli, et al. 2020]. Dystrophin gene is composed of 79 exons and has seven promotor sequences. Three of them located at the 5’ end of the gene are responsible for translation of the whole gene and synthesis 427 kDa proteins known as full dystrophins. They subtly differ in the tissue-specific manner at the N-terminus. Full dystrophin is synthesized in skeletal and smooth muscles, cardiomyocytes and neurons. Remaining four much shorter dystrophins are expressed in various tissues but none of them can replace 427 kDa. Thus lack of full dystrophin is necessary and sufficient cause of DMD. 427 kDa dystrophin is a cytoskeletal protein which joints intracellular actin filaments with proteins located in the sarcoplasmic membrane and indirectly with specific intra- and extracellular proteins. Such a multiprotein assembly is known as a dystrophin associated protein complex [Gao et al. 2015]. Understanding of these structures gave a basis for theories which pointed out a role of dystrophin in stabilization of the sarcoplasm membrane and finally muscle fibres during contraction. However, such an explanation does not apply for non-contracting cells such as neurons. In addition, lack of dystrophin has been thought to prevent proper DAP formation which is considered as a scaffold for numerous membrane proteins and an important element of the cellular signalling network [Colledge and Froehner 1998]. Yet, growing body of evidence clearly show that such a “mechanical” role of dystrophin does not explain all cellular consequences of DMD. Particularly, dystrophy-related complications in cells which do not synthesize full dystrophin are not easily explainable on a basis of this concept [Ferrari et al. 1994; For rev. see Zabłocka et al. 2021]. Myoblasts which are premature muscle cells do not contain 427 kDa dystrophin regardless of any mutations in dystrophin gene because of too early stage of their differentiation. However they exhibit many phenotypic consequences of DMD mutation [Yeung et al. 2006; Onopiuk et al. 2009; Onopiuk et al. 2015]. Although a molecular mechanism behind such a phenomenon is unknown so far, DMD consequences in cells which do not ever synthesize full dystrophin force the concepts explaining DMD pathophysiology to be reconsidered. In a course of Duchenne Muscular Dystrophy, regenerative processes may partially counteract muscle degradation as long as a pool of satellite cells is not completely exhausted, although it is not clear whether myoblasts harbouring DMD mutation may support proper regenerative processes [Chang et al. 2016. The multistep regeneration begins when satellite cells are stimulated to differentiation to myoblasts and then to myotubes and matured muscle fibres. It may be followed on a basis of changed pattern of the specific transcription factors expression. Moreover, variations of the intracellular calcium signalling that are reflected by changes in the global or local Ca^2+^ concentration are of crucial importance for muscle differentiation to occur. Calcium signalling is activated, modulated and closed due to an orchestrated action of a spectrum of molecular tools (“calcium toolkit”), which maintain appropriate distribution of Ca^2+^ and allow calcium cations to flow downhill and uphill their concentration gradient in a very precisely controlled way. Amongst the list of such tools whose expression and activity change during muscle cell differentiation and maturation nucleotide receptors have been pointed out as particularly important proteins [Ryten et al. 2004;]. They are activated by extracellular nucleotides (ATP in particular) and initiate specific cellular calcium signal. In a case of metabotropic P2RY a stimulation with nucleotide(s) cellular response involves protein G and phospholipase C while ATP-activated ionotropic P2X receptors act as a Ca2+ channels in the plasma membrane. Muscle development leading to their formation and growth is to a large extent recapitulated during muscle regeneration upon injury or in the course of muscle-degenerating diseases. Among them DMD belongs to the most prominent.

The majority of basic biochemical and molecular study on muscular dystrophy have been performed with the use of animal models. In particular *mdx* mice which do not synthesize of the full dystrophin protein due to nonsense point mutation within the exon 23 are the most commonly investigated.

Using immortalized myoblasts derived from w/t and *mdx* mice we previously showed substantial differences not only concerning nucleotide receptors but also store-operated calcium entry and many other proteins involved in the cellular calcium signalling pathways [Yeung et al. 2006; Young et al. 2012; Onopiuk et al. 2015; Róg et al. 2019]. Moreover we also pointed out *mdx*-related consequences in mitochondrial network organization and cellular energy metabolism [Onopiuk et al. 2009]. Immortalized w/t and *mdx* myoblasts had been grown in the same conditions for a long time before their use. Such a protocol allows indicating these differences which reflect intrinsic cellular consequences of DMD, not influenced by different culture conditions. However after long-term growth in artificial conditions immortalized myoblasts do not allow identifying individual properties of myoblasts derived from specific muscles. On the other hand, primary myoblasts freshly isolated from normal and dystrophic muscles may exhibit DMD-related differences not only due to their intrinsic properties but also resulted from the specific intramuscular environment. Moreover, they have tendency to spontaneous differentiation, while conditionally immortalized cells maintain their myoblast-specific properties unless grown conditions were switched to “differentiating” ones. Thus, it seems that the use of both approaches (models) will deliver more complete and compatible information. Previously we collected numerous experimental facts evidencing a very special role of P2RX7 activated by extracellular ATP in DMD pathophysiology. In contrast present study are entirely focused on the metabotropic nucleotide receptors (P2Y family) as they refer to another set of our previously published data that indicated P2RY2 as a principal metabotropic nucleotide receptor, activity of which was elevated in immortalized *mdx* myoblasts [Róg et al. 2019]. However, present data do not call into question a crucial role of ionotropic nucleotide receptors and P2X7 in particular in DMD pathophysiology. This issue has been broadly described elsewhere and is continuously investigated by our team [Yeung et al. 2006; Young et al. 2012; Young et al. 2017; Al-Khalidi et al. 2018]. Here we have tested an expression and activity of metabotropic nucleotide receptors in myoblasts obtained from four muscles isolated from w/t and *mdx* mice. Moreover, we compared a distribution of specific proteins involved in cellular calcium homeostasis in myoblasts derived from particular muscles. The spectrum of these proteins was selected on a basis of previously published results obtained for the immortalized myoblasts [Róg et al. 2019]. We have found substantial differences between myoblasts derived from various muscles of normal (dystrophin positive) mice, and shown that the consequences of *mdx* mutation exhibit muscle-specific pattern. Thus the genotype-phenotype relations are not the same in mice myoblasts isolated from various muscles. Therefore our previously obtained results with the use of immortalized *mdx* myoblasts as an experimental model of dystrophy should not be simply translated to all myoblasts irrespectively of their muscle origin, though all of them consistently confirm general thesis of aberrant calcium signalling in dystrophic cells.

## Methods

### Primary myoblasts; cell isolation and culture

Mice were used with the approval of the local ethics commission (Polish Law on the Protection of Animals). Satellite cells were prepared from hindlimb muscle of control (C57BL/10ScSnJ) and dystrophic (C57BL/10ScSn-Dmd/J) adult (8 weeks old) male mice (Jackson Laboratory, Jacksonville, USA). Isolation and purification of satellite cells from tibialis anterior (TA), Gastrocnemius (GC), soleus (Sol) and Flexor Digitorum Brevis (FDB) muscles and myoblast culture procedures were performed as described in [Musarò and Barberi L, 2010]. Briefly, whole muscles were sterilized in PBS with betadine and then digested with 0.2% collagenase type IV in DMEM for 1.5 h at 37°C, rinsed in DMEM supplemented with 1 g/l glucose, 1 mM pyruvate, 4 mM L-glutamine, 10% HS, 0.5% chicken embryo extract (CEE), 1000 U/l penicillin (1000 U/l) and 1 mg/l streptomycin and triturated with pipettes of gradually decreasing diameter. Entire fibres were separated, rinsed four times with the same medium and finally transferred into DMEM (composition as above, supplemented with 20% FBS). Purified fibres were dispersed by forcing through an 18 gauge injection needle, filtered through 40 μm pore diameter nylon bolting cloth, put into collagen-coated culture dishes and incubated in the same medium at 37°C in humidified atmosphere of air (95%) and CO2 (5%) for 3-5 days. Then, the cells were passaged and put into collagen-coated dishes prepared for the experiments.

### RNA extraction, reverse transcription and quantitative RT-PCR

Total RNA was isolated using Trizol method (TRI reagent, T9424; Sigma). The quality and quantity of samples were determined using a NanoDrop spectrophotometer. Only RNA with an absorbance ratio 260/280 between 1.8 and 2.0 was used to reverse transcription. Complementary DNA (cDNA) was synthesized from 2 μg of total RNA using First Strand cDNA Synthesis Kit with M-MLV reverse transcriptase and oligo(dT) primers (#K1612, Thermo Fischer Scientific), according to the manufacturer’s instructions. RT-qPCR was performed using TaqMan Fast Universal PCR Master Mix (4,352,042, Applied Biosystems) and TaqMan Gene Expression Assays (the primers ID: Mm00435471_m1 for p2ry1, Mm01274119_m1 for p2ry2, Mm00445136_s1 for p2ry4 and Mm01275472_m1 for p2ry6, Mm00446026_m1 for p2ry12 and Mm00546978_m1 for p2ry13) on the 7500 ABI Prism Real-Time PCR System (Applied Biosystems). The level of expression of target genes were normalised to the expression of GAPDH (primer ID: Mm99999915_g1) housekeeping gene. Analysis of relative gene expression data were determined by 2-ΔΔCt method using StepOne Software.

### Cell lysis, protein electrophoresis and analysis

Proteins were extracted from adherent cells by scraping into extraction buffer (1× LysisM, 1× protease inhibitor cocktail, 2 x phosphatase inhibitor cocktail (all Roche), 2 mM sodium orthovanadate (Sigma) suspending with the use of automatic pipets followed by repeatedly forcing through the syringe needle (0.5 mm in diameter) and incubation of the suspension on ice for 20 min. After centrifugation (15,000 ×g, for 20 min at 4 °C) protein concentrations in supernatants collected were determined using a Bradford protein assay (Bio-Rad). Remaining supernatants were mixed with sample buffer at 3:1 v /v ratio, heated for 5 min at 95 °C and chilled on ice and stored at −80 °C. Proteins (20 μg of each sample) were separated on 0.1% SDS polyacrylamide gels (6–12% w/v depending on the molecular mass of protein) and electroblotted onto Immobilon-PVDF Transfer Membrane (Merck Millipore). Blots were blocked in 5% w/v non-fat milk or 5% BSA (Albumin, Bovine Serum, 12659, Merck Millipore) powder solved in 1 × TBST, 0.01% v /v Tween-20 (Sigma) for 1 h at room temperature (RT) prior to probing with appropriate primary antibody diluted in a blocking buffer containing 2.5% milk or 5% BSA (incubated overnight at 4 °C with agitation). To identify proteins of interest following primary antibodies were applied: P2Y1 (1:270, APR-009), P2RY2 (1:270, APR-010), P2RY4 (1:300, APR-006), P2RY6 (1:250, APR-011), P2RY12 (1:270, APR-012), P2RY13 (1:270,APR-017); all Alomone Labs, calsequestrin (ab126241), calreticulin (ab128885), SERCA1 (ab124501) and SERCA2 (ab91032; all diluted 1:1000, Abcam), Gαq11 (1:1000, 06-709 Merck Millipore), PLCβ isoforms 3-4 (sc-133231, sc-166131, respectively; all diluted 1:100, Santa Cruz Biotechnology), NCX1 (1:1000, R3F1 Swant), NCX3 (1:500, ab84708 Abcam), PMCA (1:1000, ab2825 Abcam) and IP3R (diluted 1:1000 in BSA solution as described above, #8568 Cell Signalling Technology). Then membranes were washed (3×) with 1 × PBST for 10 min each wash and incubated with anti-Rabbit (diluted 1:5000, ab6721, Abcam) or anti-Mouse (1:3000, ab6728, Abcam) horseradish peroxidase-conjugated secondary antibody for 1 h at RT. Specific protein bands were visualized using luminol-based substrates (Millipore) and images obtained using a Fusion FX (Vilber Lourmat). β-tubulin (1:10000, ab21058) antibody used as a protein-loading. Densitometric analyses of specific protein bands were made using exposure times within the linear range and the integrated density measurement function of BIO-1D (Vilber Lourmat).

### [Ca^2+^]c measurements in myoblasts in vitro

Myoblasts were cultured on glass coverslips in a 35 mm culture dishes (100,000 cells) for 48 h under conditions described above. Cells (70–80% confluent) were loaded with 2 μM Fura 2 AM (Molecular Probes, Oregon) in culture medium for 20 min at 37 °C in a humidified atmosphere of 95% O2 and 5% CO2. The cells were then washed twice with the solution composed of 5 mM KCl, 1 mM MgCl2, 0.5 mM Na2HPO4, 25 mM HEPES, 130 mM NaCl, 1 mM pyruvate, 5 mM D-glucose, and 0.1 mM CaCl2, pH 7.4 and the coverslips were mounted in a cuvette containing 3 ml of either the nominally Ca2+-free assay solution (as above but 0.1 mM CaCl2 was replaced by 0.05 mM EGTA) and maintained at RT in a spectrofluorimeter (Shimadzu, RF5001PC). Fluorescence was recorded at 510 nm with excitation at 340 and 380 nm. At the end of each experiment the Fura 2 fluorescence was calibrated by addition of 33 μM ionomycin to determine maximal fluorescence followed by addition of EGTA to complete removal of Ca2+. To compare calcium content in the ER store, w/t and mdx myoblasts were treated with ionomycin in the absence of Ca2+ in the bath. Ionomycin distributes evenly through cellular membranes, thus an elevation of cytosolic Ca2+ concentration is due to a fact that its concentration between the ER and cytosol equilibrates much faster than that between the cytosol and extracellular milieu. It likely mirrors the proportion between surface area of the ER and PM membranes [35]. Though such an approach is not sufficient for calculating Ca2+ content in the ER in the absolute units, it is useful to compare a total amount of Ca2+ stored in various cells or under different conditions. Cytosolic Ca2+ concentration [Ca2+]c was calculated according to Grynkiewicz et al. [36]. The cells were treated with 500 μM ATP, ADP 1 mM, 100 μM UTP, UDP 1 mM, (all Sigma) and 10 μM AR-C 118925XX (216657-60-2, TOCRIS Bioscience) applied 10 min prior to the addition of ATP and UTP.

### Immunocytofluorescence (ICC)

Cells were cultured on coverslips (50–60%) in growth medium. After rinsing twice with cold PBS (w/o calcium and magnesium), the myoblasts were fixed in a 4% w/v paraformaldehyde solution (PFA) in PBS for 15 min on ice. Cells were then permeabilized using PBS with 0.1% Triton X-100 for 5 min and blocked in a 5% v /v goat serum (GS, Normal Goat Serum, S-1000, Vector Laboratories IVD) in PBS for 1 h at RT and incubated overnight at 4 °C with primary antibodies (aniP2RY1, antiP2RY2, anti-P2RY4 and antiP2RY6, as described above, diluted 1:100 in blocking buffer). The secondary antibody (diluted 1:1000 in 5% GS in PBS, Alexa Fluor®488 goat anti-Rabbit, Thermo Fisher Scientific) was added for 1 h RT in dark. Then myoblasts were rinsed for 10 min 3 times under agitation between each step of ICC protocol. After staining, cells on coverslips were mounted onto microscope slides sealed in Glycergel Mounting Medium with DAPI (H-1200 VectaShield®, Vector Laboratories) prior to imaging. Images were obtained using a confocal microscope (Zeiss Spinning Disk Confocal Microscope) and image analysis was performed using ImageJ software.

### Random motility assay

15,000 myoblasts (*mdx* and w/t) were seeded into 24-well cell culture plate and grown for 48 h in an appropriate culture medium. Then, multiple different well areas per each cell type were photographed in the bright field or DIC Nomarski contrast using HC PL APO 10x/0.40 Dry objective (Leica Microsystems GmbH) every 15 min for 5 h 45 min. inverted DMI6000 microscope (Leica Microsystems GmbH) equipped with DFC350FXR2 CCD camera (Leica Microsystems GmbH) and an environmental chamber (PeCon GmbH). Acquired time-lapse movies were exported to TIFF format and aligned to compensate possible drift using an ImageJ plugin. Subsequently, at least 30 cells of each experimental condition were tracked semi--+automatically in the time-lapse movies using Track Objects plug-in in Leica MM AF powered by MetaMorph^®^ software (Leica Microsystems GmbH). Cells dividing or colliding with other cells were excluded from the analysis, which has been performed as described below.

### Data analysis

Data are expressed as a mean value ± standard deviation (SD). Statistical significance was assessed by Student’s t-test A p value of < 0.05 was considered statistically significant where n = 3 – 4 for PCR, n = 3 - 6 for western blot data, and n = 3 - 5 for calcium measurements; “n” represents the number of repeated experiments with cells derived from three different mice.

## Results

Fig. 1 shows that a pattern of cellular level of transcripts encoding for various P2RYs differs in terms of both: amount of specific mRNA in myoblasts derived from particular dystrophin-positive muscles and effect of mdx mutation on the cellular content of individual receptors. Substantial differences were observed among myoblasts isolated from various muscles of w/t animals with exception of P2RY4. Also effects of mdx mutation on transcripts level do not allow bringing to the general conclusion. Comparison of P2RY4 transcript levels indicates at least strong increasing tendency in mdx myoblasts originating from all muscle tested. Also P2RY6 transcript level is substantially elevated in GC muscle and exhibits strong increasing tendency in EDB.

**Fig.1.**
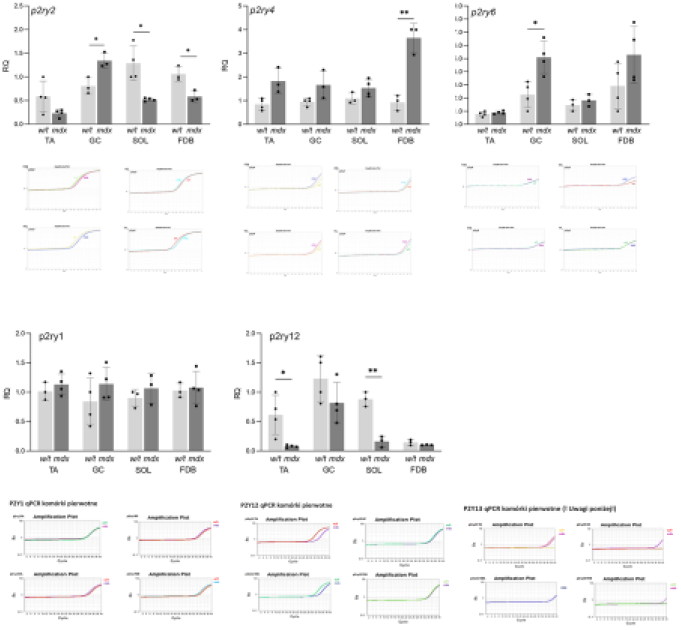
P2RYs transcripts level in myoblasts derived from four muscle. All classes of transcripts encoding P2Y2, P2Y4 and P2Y6 were tested in the same sample thus they may me compared quantitatively. Similarly mRNAs encoding three ADP-activated receptors (P2Y1, P2Y12 and P2Y13) were detected in the same samples. * p < 0.05, ** p < 0.01 (mdx vs. w/t)

**Fig. 2.**
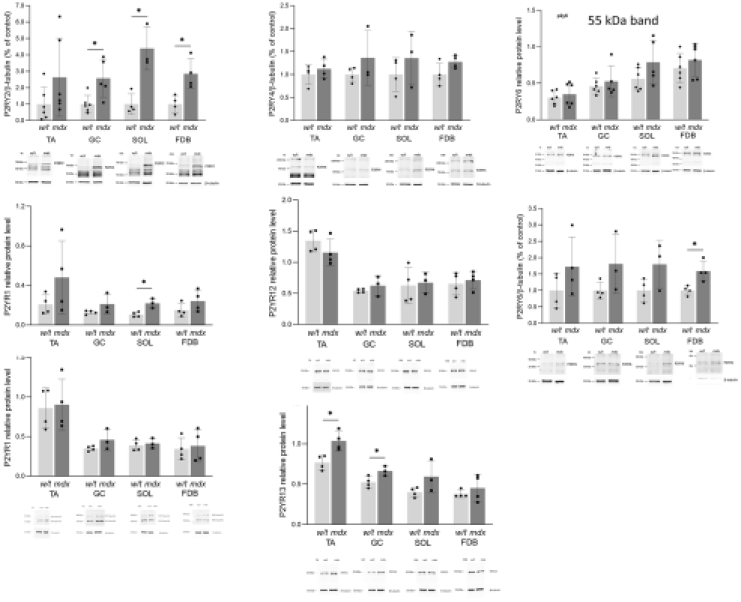
Western blots. Each Western blot shows data from randomly paired w/t and mdx mice used in one experiment. Each bar represents mean value from 3 independent experimenst ± SD. Under each set of bars representative western blots are shown. * p < 0.05, ** p < 0.005 (mdx vs. w/t)

Cellular content of mRNA for P2RY1 and P2RY12 are unaffected or declined due to *mdx* mutation in all muscle tested, respectively. More precise analysis of traces (to compare number of cycles) indicates that an expression of ADP-sensitive receptors is much lower than in a case of P2RY2 and P2YR4 and poorly detectable (P2RY13 in particular and therefore not shown).

### Western blots

In contrast to mRNA data, protein level af all 7 receptors was elevated or at least unchanged in *mdx* myoblasts isolated from all muscle tested. Particularly amount of P2RY2 was substantially increase in the myoblasts. It could suggest a special impact of P2RY2 on calcium signalling in these cells. Moreover, in vast majority of cases amount of protein of all receptors tested was similar in w/t myoblast regardles of their muscle origin. Noteworthy, *mdx* mutation never resulted in a reduced protein content in myoblasts isolated from any tested muscles.

These pictures visualise distribution of P2 receptors in myoblasts from four muscles. While a location of P2RY2 in the plasma membrane does not arouse any doubts (although quantative differences between w/t and *mdx* cells are not obvious in this matter) P2RY4, seems to localise inside cells regardless of their muscle origin. This observation is in agreement with effects of P2RY2 antagonist on calcium response (Fig. 4A). In addition in mdx myoblasts obtained from TA and DG P2RY2 seems to localise in the plasma membrane then in w/t cells while in a case of SOL and FDB any differences in this matter are invisible Nuclear staining with P2RY2 antibody reproduces effects observed in immortalized cells [Róg et al. 2019] and probably results from imperfect specificity of it. Also P2RY1 and P2Ry6 differ in their intacellular localizatio between differen muscles but an effect of *mdx* mutation on their intracellular distribution is not obvious.

**Fig. 3.**
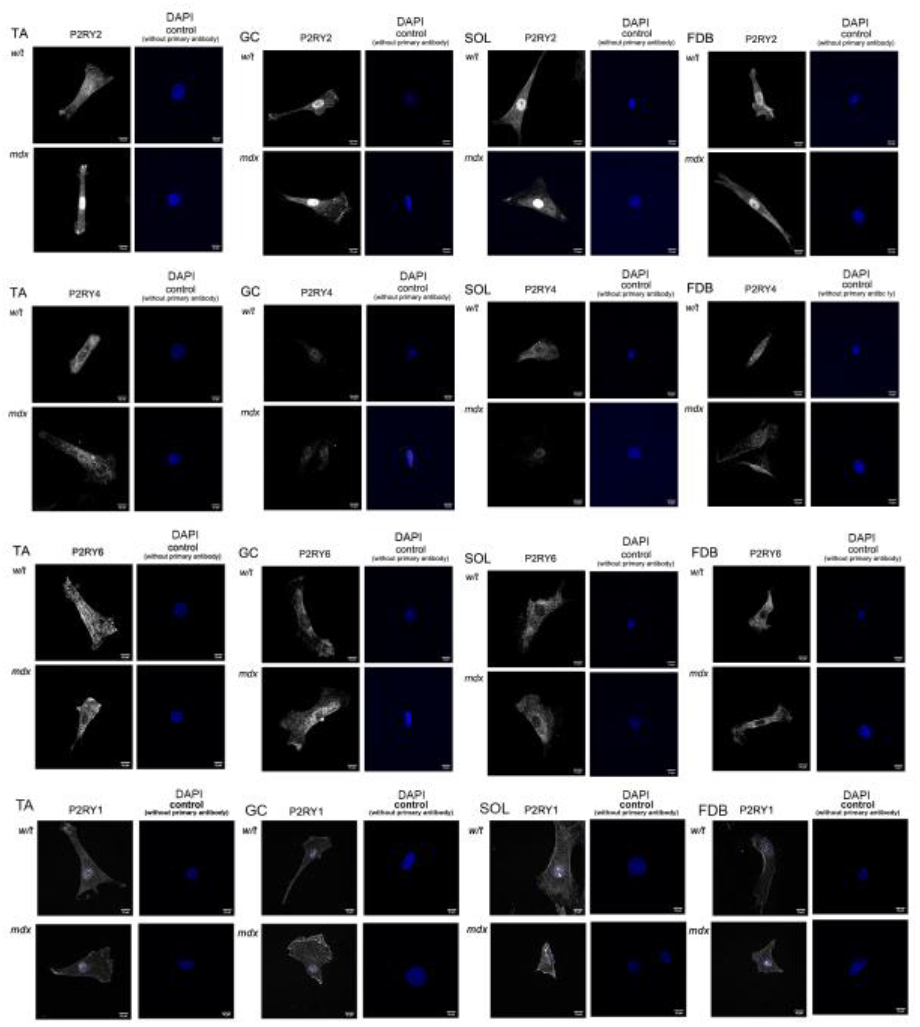
Fluorescent microscopy visualization of P2RY2, P2RY4 P2RY6 and P2RY1 in myoblasts isolated from four muscles of w/t and mdx mice. Exemplary pictures. TA, Tibialis anterior; GC, gastrocnemius; SOL, Soleus; FDB, Flexor digitorum brevis’s

**Fig. 4.**
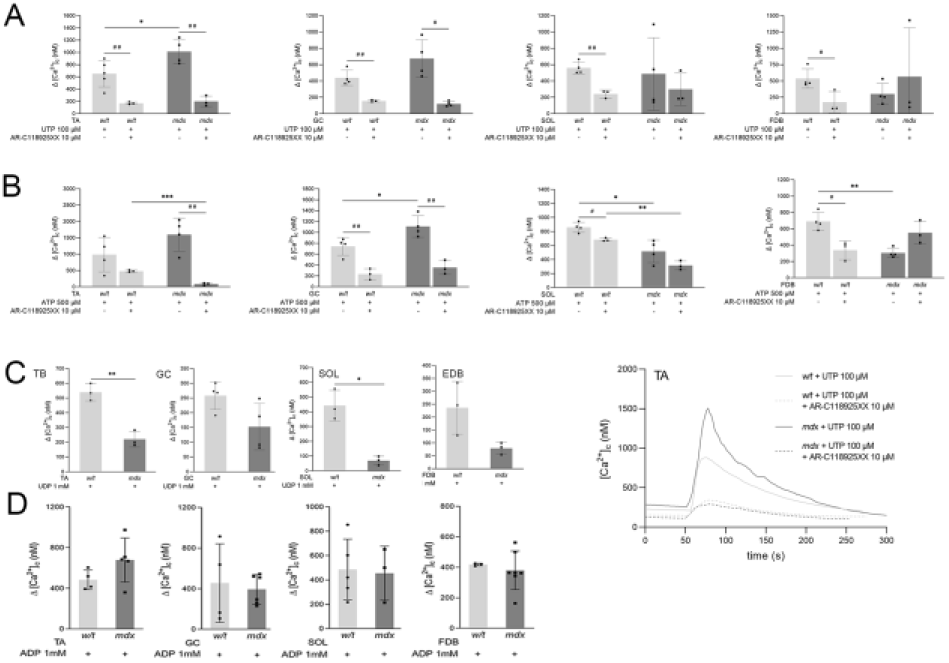
Nucleotide - induced calcium release from ER. Representative trace (bottom right) is to show a general concept of the experiment. Myoblasts derived from one out of four tested muscles were incubated with or without P2RY2 inhibitor and then 100 μM UTP (**A**) or 500 μM ATP (**B**) was added. Myoblasts derived from one out of four tested muscles were treated with UDP (**C**) or ADP (**D**). Collected data for three independent experiments. Asterix(s) * marks mdx vs. w/t and (mdx+P2RY2 antagonist) vs. (w/t + P2RY2 antagonist). Hashtag # marks mdx vs. (mdx + P2RY2 antagonist) and w/t v/s (w/t + P2RY2 antagonist). * and # p < 0.05; ** and ##p < 0.01; ***p < 0.001

### Calcium release from the ER stores

Fig. 4 shows that an intensity calcium response of w/t myoblasts, reflecting Ca^2+^-release from the ER stores due to stimulation of myoblasts incubated under Ca^2+^-free conditions (to avoid simultaneous response of P2X receptors and/or store-operated calcium entry) by specific nucleotides strongly depends on the origin of cells. Moreover, effects of *mdx* mutation on changes in cytosolic Ca^2+^ concentrations are different and not always parallel with changes concerning appropriate transcripts and proteins. The most unequivocal stimulation of Ca2+ response due to mdx mutation has been observed in myoblasts derived from TA and GC stimulated wit ATP or UTP. In contrast, a response of myoblasts from SOL or FDB seems to be opposite in the same experimental conditions. Residual calcium response observed in cells preincubated with P2RY2 antagonist prior to stimulation with ATP or UTP may reflect activity of P2RY4. Unexpectedly increased response to UTP and ATP observed in a case of *mdx* but not w/t myoblasts derived from EDB is difficult to be explained. It is noteworthy that UDP-stimulated P2RY6 activity was significantly reduced and ADP-evoked response was unaffected in all *mdx* myoblasts tested in comparison to calcium responses in their w/t equivalents. ADP is a common agonist for P2RY1, 12, 13. Because we have not tested effect of specific antagonists we cannot discriminate between them. However, our preliminary results for immortalized w/t and *mdx* myoblasts suggest that P2RY12 and P2RY13 activities overwhelm P2RY1-mediated calcium response to stimulation with ADP (Oksiejuk, unpublished data).

As it is shown in Fig. 6 a motility of the primary cells isolated from dystrophic TA and SOL was substantially more intensive than in a case of their dystrophin-positive equivalents. This observation is in line with previously published data for conditionally immortalized w/t and *mdx* myoblasts grown in the conditions that reversed immortalization (“de-immortalized cells”). In contrast to that the motility of myoblasts from GC muscle was unaffected due to *mdx* mutation. Also an inhibitory effect of ATP on dystrophic myoblasts was observed in cells derived from TA and SOL. Motility of myoblasts derived from dystrophin-positive TA and soleus was only insignificantly affected while inhibitory effect of ATP on w/t myoblasts from GC was substantial. An effect on motility of mdx cells from GC was visible but poorly significant. Although an excitation of myoblasts with ATP does not induce specific response as this agonist may not only activate metabotropic receptors P2RY2 and 4 but also ionotropic P2RX7 that was convincingly documented earlier. On the other hand however these results point out one of potentially relevant physiological consequences purinergic responses which differ dystrophic and normal myoblast. Moreover they also suggest intermuscular diversity which should be counted in experiments with myoblasts.

**Fig. 5.**
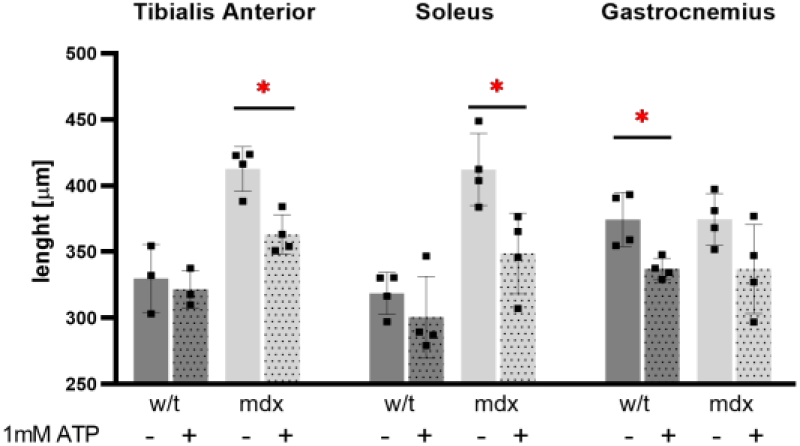
Random motility of primary myoblasts; effect of ATP-evoked cell stimulation. Collected data for 4 mice. 30 myoblasts per each muscle were tracked. * p < 0.05.

**Fig. 6.**
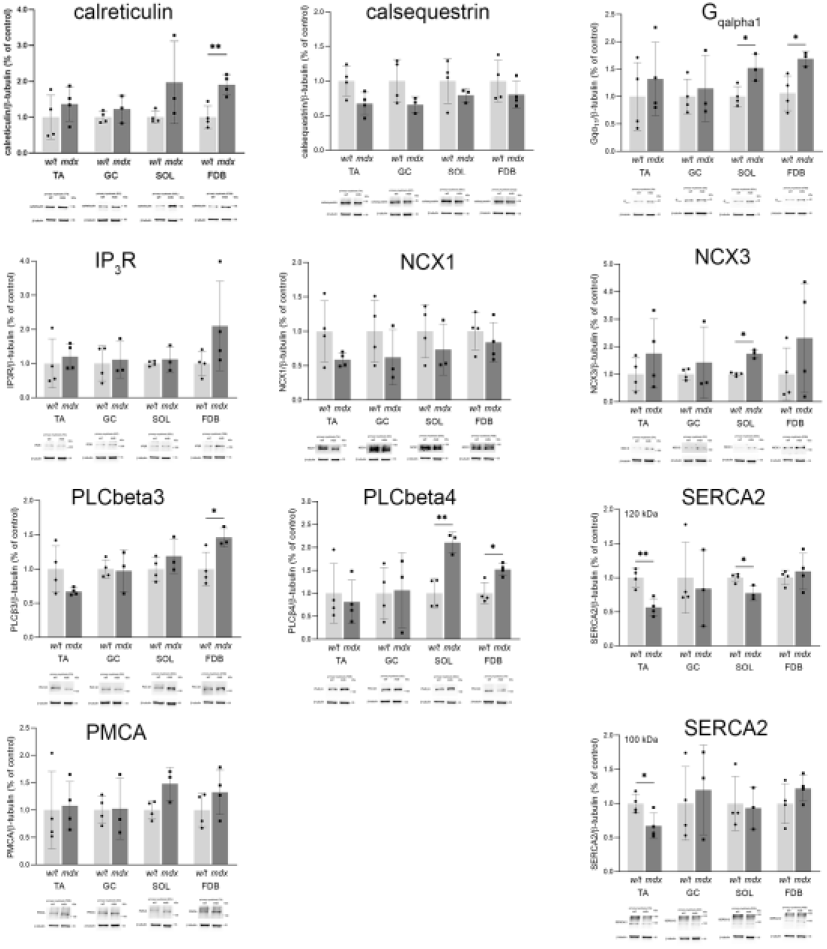
Selected proteins belonging to the calcium toolkit. Bars represent mean values for three experiments ± SD. Below each graph western blot data from one representative experiment are shown. * p < 0.005, ** p < 0.01

Data presented in Fig.6 show that *mdx* mutation influences Ca^2+^ signalling in a very complex and multifaceted means. Apart from an affected expression of the nucleotide receptors (Fig. 3) dystrophic myoblasts exhibit many other changes in the relative amounts of specific proteins belonging to the “calcium signalling toolkit” (Fig. 6). Furthermore these effects are not the same in all muscles tested. Previously we also found substantial differences between expression patterns of these proteins in immortalized w/t and *mdx* myoblasts [Róg et al. 2019]. These effects highlight a high complexity of changes in cellular calcium handling in dystrophic myoblasts in comparison to that in the w/t equivalents. It also turned out that some western blot data obtained for primary cells are less reproducible than it was in a case of immortalized myoblasts. In view of the fact that different levels of the reproducibility were observed for various proteins of the same cell lysate and visualized at the same blots it seems possible that it may reflect differences in their real impact on cellular metabolism. In other words cellular level/activity of proteins which are of particularly high regulatory function must be controlled more precisely than proteins which are less critical for global regulation of cellular metabolism, thus their amount may be controlled less tightly and fluctuate within broader range. Moreover, perfect homogeneousness (homogeneity) of primary myoblasts is more difficult to be maintained, then it is in a case of pure immortalized cultures, as they are intrinsically committed to differentiate. This also may explain relatively lower level of reproducibility.

## Discussion

Affected calcium homeostasis is a common feature of many cells harbouring mutation in the dystrophin-encoding gene, regardless of the fact whether full dystrophin is normally synthesized in these cells or not [Zablocka et al. 2021]. Our earlier study led us to conclusion that myoblasts with *mdx* mutation are more susceptible to ATP stimulation mainly because of increased activity of P2RY2 and P2RX7. Remaining nucleotide receptors seem to be less important in this context. It looks possible that both metabotropic and ionotropic ATP-sensitive nucleotide receptors act compatible upon excitation of cells by their common stimulus. Moreover, abnormally intensive calcium response of *mdx* myoblasts to ATP result from substantial changes in a broad spectrum of other “calcium toolkit” elements such as calcium pumps, exchangers and buffers [Róg et al. 2019; Zablocka et al. 2021]. The present study aim at testing whether experimental data obtained previously for cells which were substantially modified to accomplish their immortalization may be translated to primary cells reflecting more physiological situation. Furthermore, immortalized cells could not be attributed to specific myoblasts derived from any particular muscle. To address the problem of muscle specific properties of w/t myoblasts and effects of *mdx* mutation on an expression and activity selected proteins we have examined primary myoblasts derived from TG, GC, Soleus and FDB muscles. In these study we have narrowed our interest to metabotropic receptors although effects of *mdx* mutation on P2RXs expression and activity in primary myoblasts isolated from various muscles were also confirmed by our team (not shown, to be published elsewhere). All experimental results presented here may be analysed comparatively according to a few criterions. For example each receptor may be discussed separately in all muscles or all receptors in one selected muscle. Both assessments clearly indicate that the pattern of P2RYs distribution and the level of specific transcripts among myoblasts derived from four muscles tested differs. Moreover, effects of *mdx* mutation on P2RYs expression and activity is also diverse and not always parallel with changes in receptor protein level. The origin of these differences is not clear but it may be speculated that they reflect specific metabolic features of particular muscles and specific changes in the local “dystrophic environment”. It may diversely influence myoblast gene transcription. Thus an epigenetic factor could also be considered. However small spectrum of muscles presented here does not allow us to convincingly discuss a putative relation between metabolic profiles of particular muscle (slow-switch vs. fast-switch) and its sensitivity to *mdx* mutation particularly in a context of nucleotide receptors presence and activity. Undoubtedly this problem is worth addressing in the future.

The most evident *mdx*-induced changes within P2Y receptor family concern P2RY2 protein level that is increased in myoblasts derived from all muscle tested. Moreover, this is the only P2RY of which activity is elevated (not always with high significance) in *mdx* myoblasts regardless of muscle of their origin. These observations strongly resemble our previously described data obtained with the use of immortalized myoblasts derived from w/t and *mdx* mice [Róg et al. 2019] and strengthens their physiological reliability. On the other hand however, the pattern of distribution of other proteins belonging to “calcium toolkit” differ not only among primary myoblasts used in the present study (see Fig. 5) but also does not fit in with the data found for immortalized cells [Róg et al. 2019]. Thus it seems that immortalized myoblast we used previously may be considered as a valuable model for basic biochemical study which emphasize intrinsic properties distinguishing w/t and mdx cells. However, more “physiological” conclusions must be drawn carefully with the consciousness that both immortalization procedure as well as taking of cells out of tissue context may induce poorly predictable side effects. Immortalized cells used in our earlier experiments as well as commonly used C2C12 cells have undefined muscular origin and/or potential “loss of the memory of origin” concerning the specificity of particular muscle that they were isolated from. On the other hand, the very procedure of isolation of primary cells from specific mice muscles which is inseparable from an oxidative stress may also have an uncontrolled impact on quality of cells. Therefore, both experimental approaches are useful and complementary to each other but none of them is free from imperfections.

While a comparison of particular protein levels among myoblasts derived from different muscles or different animals (w/t and *mdx*) using Western blot should be interpreted with caution as this method of protein detection is only semi-quantitative, the levels of particular transcripts determined with the use of qPCR are more reliable. On the basis of numbers of cycles of TaqMan PCR assays which are shown shown in Fig 1 we suggest the only truly important effects of *mdx* was found in a case of P2RY2 and P2RY4 as transcripts of the remaining receptors were hardly detectable. Our results clearly indicate relative variability among myoblasts derived from different muscles in terms of P2RYs expression and activity as well as a distribution of selected protein belonging to the cellular calcium toolkits. This finding shows that any effects observed on immortalized cells or primary myoblasts isolated from defined muscle should not be generalized and translated directly to myoblasts originating from any other. Results presented hitherto have only a descriptive value but considering a common use of myoblasts in a variety of projects and study it seems to be important to point out that some properties of these cells are not universal among myoblasts as a whole. As shown in Fig 5, some proteins were identified with high reproducibility while in a case of other we found bigger differences between experiments. Because all proteins were detected in the same lysates, poor reproducibility in a case of only some of them may suggest that their cellular level/activity does not have to be rigorously controlled and enforced, thus some flexibility has not been eliminated. In other words cellular level/activity of proteins which are of particularly high regulatory function must be controlled more precisely than proteins which are less critical for global regulation of cellular metabolism, thus their amount may be controlled less tightly. Moreover, perfect homogeneity of primary cells is more difficult to be maintained in comparison to highly pure immortalized cultures as they exhibit high tendency to enter differentiation process. This also may explain lower level of reproducibility.

Using primary myoblasts originating from various muscle we have confirmed previously described abnormalities concerning calcium signalling that were found in immortalized dystrophic myoblasts. Moreover we proved our earlier finding that stimulation of P2RY2 slows down a motility of dystrophic myoblasts while w/t cells are more resistant in this matter. Actually, ATP may stimulate not only P2RYs but also P2RXs, thus, the data presented here do not allow us to point out specific ATP-sensitive receptor(s) behind these differences. Particularly that our preliminary data convincingly show substantial differences in P2RX7 expression and activity in myoblasts obtained from various muscles of normal and mdx mice (data not shown, in preparation). On the other results presented her they underline a physiological context of changes in ATP-induced response of normal and dystrophic myoblasts.

Finally all od results presented here strengthen a concept of very early, in terms of muscle differentiation, biochemical consequences of Duchenne type dystrophy, at least in the mice model. Moreover they stress the fact that myoblasts derived from various mice muscles exhibit similar but not identical pattern of changes related to intracellular calcium signalling. All of these results together with our previously described effects of dystrophy on P2RX7 – mediated ionotropic nucleotide response of *mdx* myoblasts enrich and broaden the knowledge about DMD pathophysiology.

## FUNDING INFORMATION

This work was supported by the National Science Centre Poland, grant number 2013/11/B/NZ3/01573

## Notes

### Competing Interest Statement

The authors have declared no competing interest.

### Summary of Updates

Funding information

